# Low predictive power of clinical features for relapse prediction after antidepressant discontinuation in a naturalistic setting

**DOI:** 10.1101/2020.01.28.922500

**Authors:** Isabel M. Berwian, Julia G. Wenzel, Leonie Kuehn, Inga Schnuerer, Erich Seifritz, Klaas E. Stephan, Henrik Walter, Quentin J. M. Huys

**Affiliations:** Translational Neuromodeling Unit, University of Zurich and ETH Zurich, Zurich, Switzerland; Hospital of Psychiatry, University of Zurich, Zurich, Switzerland; Charité Universitätsmedizin, Campus Charité Mitte, Berlin, Germany; Wellcome Centre for Human Neuroimaging, University College London, London, UK; Max Planck Institute for Metabolism Research, Cologne, Germany; Division of Psychiatry and Max Planck Centre for Computational Psychiatry and Ageing Research, University College London, London, UK; Camden and Islington NHS Foundation Trust, London, United Kingdom

## Abstract

1

**Background:** The risk of relapse after antidepressant medication (ADM) discontinuation is high. Predictors of relapse could guide clinical decision-making, but are yet to be established.

**Method:** We assessed demographic and clinical variables in a longitudinal observational study before antidepressant discontinuation. State-dependent variables were re-assessed either after discontinuation or before discontinuation after a waiting period. Relapse was assessed during six months after discontinuation. We applied logistic general linear models in combination with least absolute shrinkage and selection operator and elastic nets to avoid overfitting in order to identify predictors of relapse and estimated their generalisability using cross-validation.

**Results:** The final sample included 104 patients (age: 34.86 (11.1), 77% female) and 57 healthy controls (age: 34.12 (10.6), 70% female). 36% of the patients experienced a relapse. Treatment by a general practitioner increased the risk of relapse. Although within-sample statistical analyses suggested reasonable sensitivity and specificity, out-of-sample prediction of relapse was at chance level. Residual symptoms increased with discontinuation, but did not relate to relapse.

**Conclusion and Relevance:** Demographic and standard clinical variables appear to carry little predictive power and therefore are of limited use for patients and clinicians in guiding clinical decision-making.

## 2 Introduction

Depressive disorders are a major burden to societies worldwide, being the single largest contributors to years lived with disability (WHO, 2017). This is largely due to the often chronic relapsing nature of depression, which underlies the functional and social impairments it brings (Lépine and Briley, 2011). Hence, successful treatment of a particular depressive episode, e.g. by achieving response to an antidepressant medication (ADM), is critical, but only the first step.

Thus, the prevention of relapses is the next step as over half of the patients with one depressive episode will experience a second one and the risk of relapse only increases further thereafter (American Psychiatric Association, 2000). Preventing relapses is of paramount importance for the longer-term course of the illness, and a number of strategies exist. One important strategy is continuation and maintenance treatment with ADM, which reduces the risk of relapse (Geddes *et al*., 2003; Kaymaz *et al*., 2008; Glue *et al*., 2010; Sim *et al*., 2015). However, there is still a risk of breakthrough depression, i.e. the development of further depressive episodes while taking ADMs (Rush *et al*., 2006). Furthermore, many patients want to discontinue their ADM due to side-effects such as weight gain and sexual dysfunction (Olfson *et al*., 2006) or adhere only partially (Hunot *et al*., 2007). At the same time, one in three patients relapses within six months after discontinuation (Geddes *et al*., 2003).

Thus, not all patients benefit equally from continuation treatment and there appears to be variation in individual trajectories after the initial response to ADMs (Uher *et al*., 2010; Gueorguieva *et al*., 2011; Muthén *et al*., 2011; Musliner *et al*., 2016). Markers that identify these trajectories and separate those patients who can safely discontinue their ADMs from those with a higher risk of relapse after discontinuation clearly have the potential of improving this situation.

Indeed, current guidelines take some of this variation into account, and recommend continuation treatment for four to nine months after the first depressive episode and two years or more after recurrent episodes (Bauer *et al*., 2013; NICE, 2010). More recently, guidelines also refer to residual symptoms and physical and psychological comorbidities (NICE, 2019). These recommendations are based on evidence derived from the natural course of depression and overall relapse risk (Frank *et al*., 1991; Kessler *et al*., 2003; Nierenberg *et al*., 2010). However, the importance of these markers has been disputed (Berwian *et al*., 2017): two meta-analyses came to diametrically opposed conclusions about the relevance of the number of prior episodes (Viguera *et al*., 1998; Kaymaz *et al*., 2008), and five separate meta-analyses have failed to find an effect of length of ADM treatment on relapse risk after discontinuation (Viguera *et al*., 1998; Geddes *et al*., 2003; Kaymaz *et al*., 2008; Glue *et al*., 2010; Andrews *et al*., 2011).

Several other predictors of relapse after discontinuation exist in the literature (Berwian *et al*., 2017). These include ethnicity (Trinh *et al*., 2011), neurovegetative symptoms (McGrath *et al*., 2000), melancholic subtype (McGrath *et al*., 2000), anxiety (Joliat *et al*., 2004), somatic pain (Fava *et al*., 2009) and response pattern to drug (Stewart *et al*., 1998; Nierenberg *et al*., 2004). Only the last predictor, having a placebo drug response, i.e. fast but unstable response, compared to a true drug response, i.e. slower but sustained response (Quitkin *et al*., 1987), has been replicated (Stewart *et al*., 1998; Nierenberg *et al*., 2004). Unfortunately, assessing this measure will be difficult in clinical practice.

Two further points complicate this picture. The first point relates to a methodological problem. Studies have usually mainly focused on asking whether a particular variable differs between groups of patients who do and do not go on to relapse. Unfortunately, while such differences might reach statistical significance, they might still fail to perform well as predictors (Lo *et al*., 2015). Regressions analyses have been used in some studies, but using a simple regression bears the risk of overfitting (Huys *et al*., 2016). To make inferences about a new patient in a practice, the predictive power in cases outside of the sample used to fit the regression model needs to be determined, e.g., using cross-validation. To our knowledge, this has not been reported in the literature so far. Second, most studies have been performed in the setting of double-blind RCTs. While this is the ideal approach to examine whether an active compound has a causal role in reducing relapse, it may underestimate relapse rates after discontinuation because medication discontinuation might have psychological effects in addition to direct pharmacological effects.

Here, we report findings from the AIDA study - a two-centre, longitudinal, naturalistic observational study of antide-pressant discontinuation. Our first aim was to investigate the extent to which variables which are easily assessable in a naturalistic setting can predict individual relapse risk and possibly guide the decision to discontinue or not. We paid specific attention to previously reported clinical predictors and examined their performance in a naturalistic setting. A secondary goal of this study was to understand the effects of discontinuation itself and how these relate to relapse. Accordingly, we investigated if any of the state-dependent variables changed with discontinuation and if that change differed between relapsers and non-relapsers.

## 3 Methods and Material

### 3.1 Participants

We recruited patients who decided to discontinue their medication independently from study participation after they were diagnosed with Major Depressive Disorder (MDD) and had a) experienced one severe (Wakefield and Schmitz, 2013) or multiple depressive episodes, b) initiated antidepressant treatment during the last depressive episode and c) now achieved stable remission, i.e. a score of less than 7 on the Hamilton Depression Rating Scale 17 (Williams, 1988) for 30 days. To identify disease and medication effects, we also recruited healthy controls (HC) matched for age, sex and education. See section S1.1 for detailed inclusion and exclusion criteria. All participants gave informed written consent and received monetary compensation for the time of participation. Ethical approval for the study was obtained from the cantonal ethics commission Zurich (BASEC: PB_2016-0.01032; KEK-ZH: 2014-0355) and the ethics commission at the Campus Charité-Mitte (EA 1/142/14), and procedures were in accordance with the Declaration of Helsinki.

### 3.2 Study Design

The study design is depicted in Fig. 1. Trained staff interviewed remitted patients on ADM to assess in- and exclusion criteria during a baseline assessment (BA). The BA consisted of the assessment of current symptoms and present and past diagnoses, as well as a short neuropsychological testing and a questionnaire batch assessing stable traits. Patients meeting inclusion criteria were randomised to one of two study arms. Of note, the first 10 participants at each site were all assigned to arm 1W2 (1/2 represents the number of the main assessment, “W” represents withdrawal). Participants in arm 1W2 underwent the first main assessment (MA1) including a questionnaire assessing state variables, then gradually discontinued their medication over up to 18 weeks and then underwent a second main assessment (MA2). Participants in arm 12W underwent both main assessments before withdrawal. This two-arm design allowed us to identify discontinuation effects while controlling for time, learning and repetition effects. After discontinuation, all patients entered a follow-up period of six months. During that period, they were contacted for telephone assessments at weeks 1, 2, 4, 6, 8, 12, 16 and 21 to assess relapse status. If telephone assessment indicated a possible relapse, patients were invited to an on-site structured clinical interview (SCID-I (Wittchen and Fydrich, 1997)) to assess criteria for relapse, i.e. fulfilling the diagnosis of a depressive episode according to the Diagnostic and Statistical Manual of Mental Disorders (4th ed., text rev.; DSM-IV-TR; American Psychiatric Association, 2000). If these criteria were fulfilled, they underwent a final assessment (FA). If no relapse occurred, the FA took place in week 26. HC underwent MA1 only. In addition to the measures reported here, participants also underwent functional magnetic resonance imaging, a range of behavioural task, electroencephalography and blood sampling during the main assessments. See supplementary section S1.2 for detailed procedures of the assessment sessions and S1.3 for observer-rated and self-report measures. Participant recruitment took place between July 2015 and January 2018.

**Figure 1:**
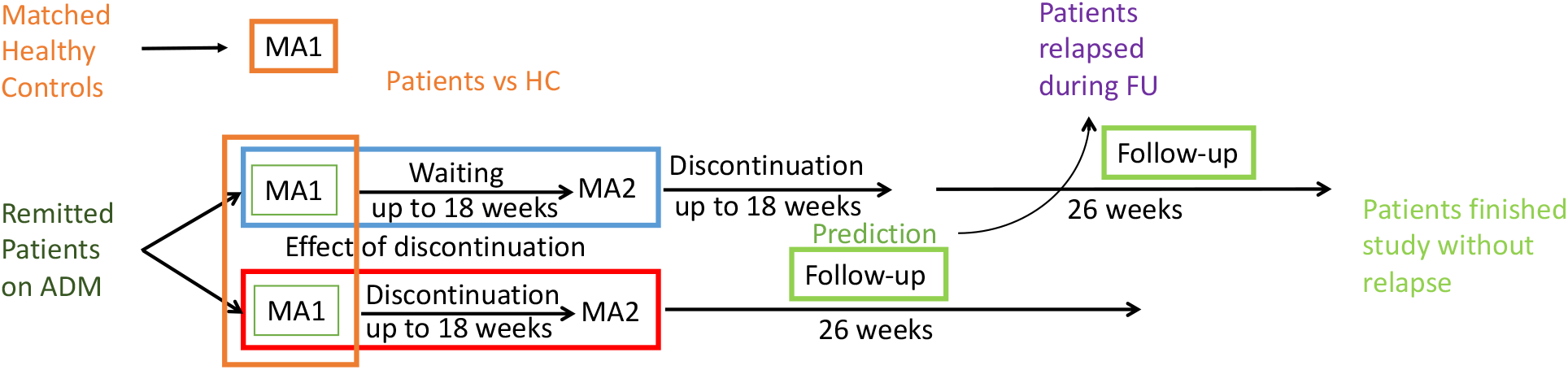
Study Design: We recruited remitted, medicated patients on antidepressant medication (ADM) and matched healthy controls (HC). They were assessed and compared at main assessment 1 (MA1) to identify traits characterising the remitted, medicated state. Next, patients were randomised to either discontinue their medication before MA2 (bottom arm, “discontinuation group” or enter a waiting period matched to the length of discontinuation time (top arm, “waiting group”). Differences in changes between MA1 and MA2 in the two separate groups were investigated to gain an understanding of the effects underlying discontinuation. Patients in the waiting group discontinued their ADM after MA2. After discontinuation, all patients entered the follow-up (FU) period of six months, whereas some patients had a relapse during this period and some patients finished this period without relapse. Differences in characteristics at MA1 of patients who relapsed and patients who did not relapse during FU provide information on which variables relates to relapse risk and can be used to identify predictors of relapse after ADM discontinuation.

### 3.3 Measures

We included 18 measures spanning four categories: demographics, current symptoms, clinical history and treatment. Measures were chosen based on two criteria: 1) they have previously been related to relapse after antidepressant discontinuation(Berwian *et al*., 2017), and 2) they can easily be assessed during a routine clinical visit, do not require extensive training or equipment and have a plausible relation to relapse risk. Individual measures in each category are listed in Table 1 and described in supplementary section S1.3. Ten of these variables were previously investigated in randomised controlled trials (RCTs). All listed measures can be assessed before discontinuation and will be included in the prediction analysis. We additionally compared discontinuation time between relapsers and non-relapsers, but did not include it in the prediction model. All measures from the category residual symptoms were re-assessed at MA2.

**Table 1:** Participant characteristics and complete-case analyses

|  | Patients vs. HC |  |  | Relapse vs. No relapse |  |  | full GLM |  | regularised GLM |
| --- | --- | --- | --- | --- | --- | --- | --- | --- | --- |
|  | Patients (n = 104) | HC (n = 57) | p value | Relapse (n = 30) | No relapse (n = 54) | P Value | coefficients | p value | coefficients |
| **Demographics** |  |  |  |  |  |  |  |  |  |
| Age | 34.85 (11.10) | 34.12 (10.95) | 0.69 | 37.00 (10.95) | 33.56 (11.23) | 0.18 | 0.073 | 0.91 | 0 |
| Male sex, No. (%) | 24 (23) | 17 (30) | 0.35 | 7 (23) | 13 (24) | 0.94 | -0.08 | 0.77 | 0 |
| Intelligence<sup>b</sup> | 28.12 (4.43) | 27.79 (4.13) | 0.65 | 29.05 (3.76) | 27.8 (4.64) | 0.09 | -0.38 | 0.31 | -0.18 |
| Site Berlin, No. (%) | 28 (27) | 22 (39) | 0.13 | 9 (30) | 14 (35) | 0.69 | 0.4426 | 0.20 | 0 |
| **Clinical predictors** |  |  |  |  |  |  |  |  |  |
| *Current symptoms* |  |  |  |  |  |  |  |  |  |
| Residual depression<sup>b</sup> | 3.73 (3.82) | 0.77 (1.25) | <0.001 | 3.77 (5.23) | 3.15 (2.59) | 0.47 | 0.15 | 0.65 | 0 |
| Residual anxiety<sup>b</sup> | 2.89 (2.34) | 1.63 (1.76) | <0.001 | 3.03 (3.02) | 2.56 (2.01) | 0.39 | 0.61 | 0.20 | 0 |
| Somatic pain<sup>b</sup> | 0.32 (0.23) | 0.15 (0.26) | <0.001 | 0.39 (0.23) | 0.28 (0.22) | 0.042 | -0.63 | 0.08 | -0.27 |
| General impairment<sup>b</sup> | 0.31 (0.25) | 0.12 (0.18) | <0.001 | 0.35 (0.31) | 0.25 (0.19) | 0.07 | -0.87 | 0.13 | -0.06 |
| *Clinical history* |  |  |  |  |  |  |  |  |  |
| Age of onset | - | - | - | 25.00 (9.88) | 23.8 (8.34) | 0.56 | 0.07 | 0.91 | 0 |
| Chronicity<sup>b</sup> | - | - | - | 8.07 (10.44) | 8.15 (9.42) | 0.97 | 0.35 | 0.30 | 0 |
| Severity<sup>b</sup> | - | - | - | 6.97 (1.13) | 7.04 (1.30) | 0.80 | 0.19 | 0.56 | 0 |
| Number of prior episodes | - | - | - | 2.77 (1.79) | 2.28 (1.48) | 0.18 | -0.42 | 0.58 | 0 |
| Severity factor<sup>c</sup> | - | - | - | 0.09 (0.38) | -0.03 (0.33) | 0.15 | 0.20 | 0.84 | -0.32 |
| Comorbidities<sup>b</sup> | - | - | - | 0.70 (1.02) | 0.80 (1.17) | 0.71 | 0.08 | 0.81 | 0 |
| *Treatment* |  |  |  |  |  |  |  |  |  |
| Treated by GP, No. (%) | - | - | - | 10 (33) | 8 (15) | 0.048 | -0.94 | 0.0051 | -0.29 |
| Duration of ADM intake<sup>c</sup> | - | - | - | 24 (29) | 22.5 (38) | 0.66 | -0.13 | 0.72 | 0 |
| Medication load<sup>c</sup> | - | - | - | 0.0068 (0.0041) | 0.008 (0.004) | 0.26 | 0.15 | 0.62 | 0 |
| Psychotherapy<sup>c</sup> | - | - | - | 0.38 (0.39) | 0.39 (0.40) | 0.95 | -0.54 | 0.12 | 0 |
| Tapering in days | - | - | - | 51.10 (40.64) | 48.89 (39.79) | 0.81 | - | - | - |

a) Unless stated otherwise, mean (SD) are shown; b) Determined as follows: intelligence: Mehrfachwahl Wortschatz Test (Lehr, 2005); residual depression: Inventory of Depressive Symptomatology-Clinician Rated (Rush *et al.*, 1996); residual anxiety: screening generalised anxiety disorder (GAD-7; Spitzer *et al.*, 2006); somatic pain: somatisation subscale of symptom checklist 90 (SCL-90; Derogatis and Cleary, 1977); general impairment: global severity index of SCL-90; chronicity: numbers of months sick within the last 5 years; severity: symptoms during the last episode; comorbidities: number of past and present psychiatric diagnoses; c) Computation of the variables is described in the section S1.4; GP = general practitioner; ADM = antidepressant medication; HC = healthy controls; GLM = general linear model. Intercept for regression models not shown.

### 3.4 Data Analysis

Analyses were performed using Matlab version 9.1.0.441655 (R2016b) according to a pre-specified analysis plan.

#### 3.4.1 Association Analyses

Candidate predictor variables were first identified by assessing group differences between patients and HC and between relapsers and non-relapsers. Two-sample two-tailed independent t-tests were used for continuous and chi-squared tests for categorical variables (including psychotherapy). We report results using no multiple comparison correction, i.e. considering tests to be significant at p < 0.05 and indicate if they survive correction using false discovery rate (FDR). The former allows for better interpretation of non-significant findings. The latter helps to control for the number of tests we applied since we are investigating a range of variables increasing the risk of false positives. In contrast to Bonferroni correction, FDR-based corrections do not make the assumption that tests are independent.

Complete-case analyses can yield biased results. We therefore examined whether patients who dropped out differed from patients who finished the study. For this, we repeated the above analyses procedure comparing patients who finished the study and patients who dropped out after MA1. We next performed Cox proportional hazards regression models, relating predictor variables to time to relapse or dropout. For these analyses, all variables were mean-centered and normalised. We first performed this for each measure individually and then included all measures in the same Cox regression, to compare predictors. Since our goal is to predict relapse after antidepressant discontinuation, we performed the latter analaysis first for the time after discontinuation, but repeated the analysis by extending the observation period to include the time of discontinuation.

#### 3.4.2 Prediction Analyses

To examine whether clinical variables have predictive value, we first fitted a full logistic general linear model (GLM) including all relapsers and non-relapsers to determine which variables made a significant contribution to the prediction, the total variance that can be explained by the combined predictors, the area under the curve, the best threshold as well as the sensitivity and specificity at this threshold.

However, as there are 18 predictors for 84 data points any results for the current sample may generalise poorly due to overfitting. To address the high number of predictors compared to the small sample size, we used an elastic net with both an L1 and L2 regularisation (Zou and Hastie, 2005) as implemented by the lassoglm function in Matlab. We applied tenfold cross validation with stratification to optimize strength of the L1-norm (λ). This was repeated for a range of *α* values and the optimum was chosen.

Next we repeated this entire procedure within a nested cross-validation procedure to examine generalisation to data not seen by the algorithm. The outer loop consisted of a leave-one-out cross-validation (LOOCV). One subject was first set aside, then the full GLM or the regularised GLM, respectively, was fitted to all other subjects. Then, the group membership of the left-out subject was predicted using parameter estimates (regression weights and optimised threshold) obtained from the other subjects. In those cases where all regression coefficients were shrunk to 0 the classification threshold was set to 0.5. These predictions were used to compute the balanced accuracy and the probability that these predictions would not be better than chance was determined with a binomial test. To determine receiver operating curves for left out subjects, we categorised these subjects as relapsers or non-relapsers for varying thresholds and computed how many subjects were categorised correctly for each threshold.

#### 3.4.3 Discontinuation analyses

To investigate the discontinuation effect and the interaction between discontinuation and relapse, we applied mixed analyses of variance (ANOVAs) with group (1W2 vs. 12W) and (relapse vs. no relapse in the discontinuation group, i.e. patients who discontinued before MA2, only) as between-subjects factor and time (MA1 vs. MA2) as within-subject factor.

## 4 Results

### 4.1 Participants

Nineteen (15%) of 123 included patients dropped out of the study prior to the first main assessment and were not further analysed. Of the 104 who completed the first main assessment, 91 (88%) completed both main assessments (44 off medication after discontinuation in arm 1W2 and 47 on medication prior to discontinuation in arm 12W). Of these, 89 (86%) achieved antidepressant discontinuation and 83 (67%) reached a study endpoint by either remaining in remission for 6 months, or only restarting antidepressants after reaching criteria for relapse. One additional patient was categorised as relapser after meeting criteria for relapse for 10 days (shorter than the length criterion of 14 days) and quick improvement after treatment re-initiation. Detailed reasons for dropouts are depicted in Figure S1.

### 4.2 Association analyses

#### 4.2.1 Complete-case analysis

Patients and healthy controls (n=57) were matched for demographic variables but patients had elevated residual depression (t(159)=5.68, p<0.001, CI=1.93-3.99), anxiety (t(159)=3.56, p<0.001, CI=0.56-1.96) and somatic pain symptoms (t(159)=4.47, p<0.001, CI=0.098-0.254) and scored higher on general impairment (t(159)=5.02, p<0.001, CI=0.11-0.26; Table 1). These results survived correction for multiple comparison.

We first performed a complete-case analysis on the 84 patients who either reached the follow-up period with-out relapse or relapsed during that period to maximise the chances of identifying potentially predictive variables. Patients who went on to relapse after ADM discontinuation had increased somatic pain (t(82)=2.07, p=0.042, CI=0.004-0.21) and were more often treated by a general practitioner rather than a psychiatrist (*χ*^2^ = 3.93, p=0.048), though these differences did not survive correction for multiple comparisons (Table 1). To assess the unique contributions of the predictor variables, all measures were combined in a single multiple regression model. This revealed treatment by GP as the sole significant variable associated with relapse (b = −0.94, p = 0.005; Tab. 1).

Complete-case analyses may yield biased results. Patients who dropped out had more residual symptoms (t(102)=-2.01, CI:-3.73– −0.025, p=0.047) and more symptoms during the last episode (t(102)=-2.09, CI:-1.24– −0.033, p=0.039) (Table S1). These differences did not survive correction for multiple comparisons.

#### 4.2.2 Intention-to-treat analyses

As there were differences between patients who completed the study and those who dropped out, we performed intention-to-treat analyses using Cox proportional hazards including all the 89 patients who completed discontinuation. This revealed that general impairment (b=0.32, p=0.044, CI=0.008-0.632) and treatment by GP (b=0.36, p=0.025, CI=0.045-0.666) were significantly associated with shorter time to relapse, though neither survived correction for multiple comparisons. Of note, no effect was found for the current symptoms and symptoms during the last episode which distinguished patients who dropped out (Table 2). To assess the unique contributions of the predictor variables, all measures were combined in a Cox multiple regression model. This again revealed treatment by GP as the uniquely significant predictor (b = 0.662, p = 0.005; Table 2, Figure 2A). GP treatment was also the only variable associated with shorter time to relapse in an extended intention-to-treat analysis including an additional 6 patients who initiated but did not complete antidepressant discontinuation (Table 2).

**Figure 2:**
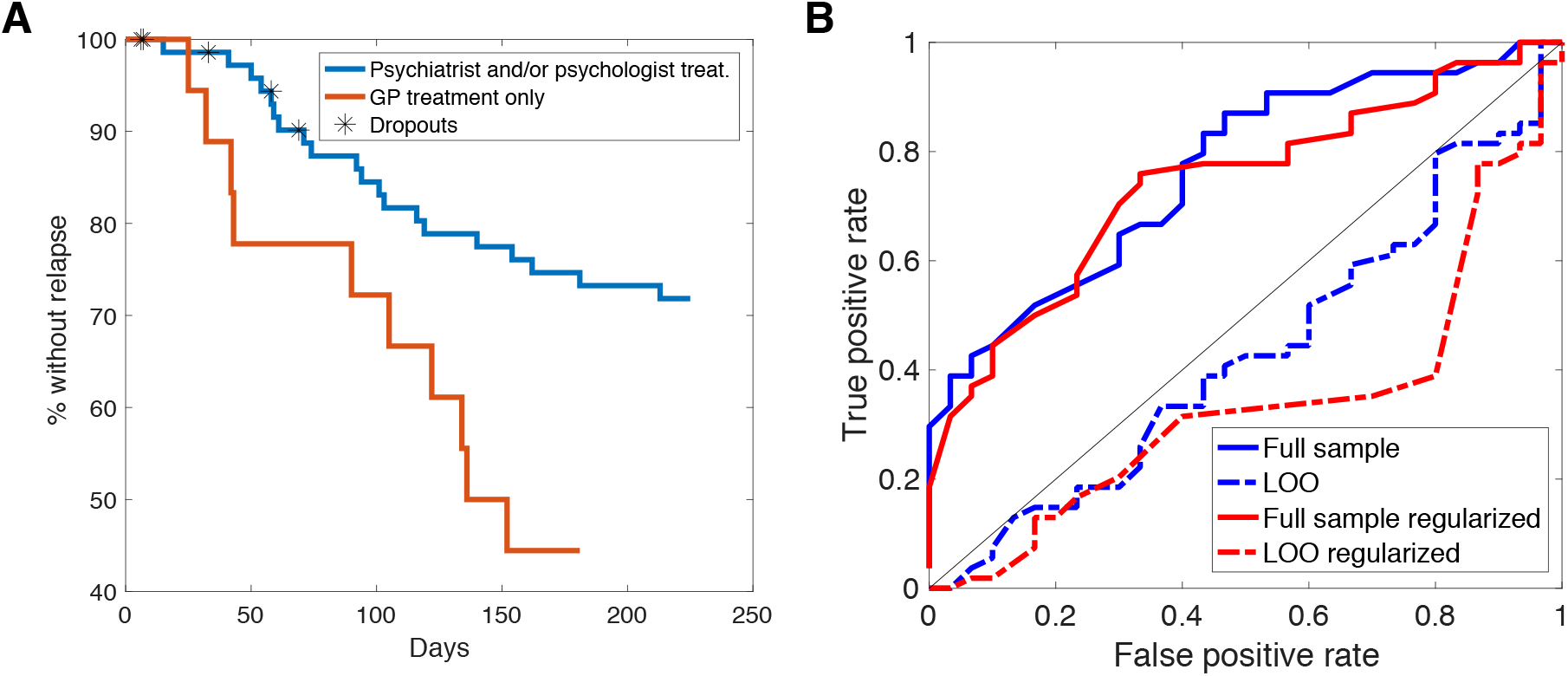
(A) Survival curves for time until relapse during follow-up period for patients who were only treated by a general practitioner (GP) or additionally by a psychiatrist or psychologist. (B) Prediction: Receiver operating curves for a standard general linear model (blue) and a regularised general linear model using least absolute shrinkage and selection operator and elastic net (red) using the full sample (solid lines) and for subjects left out of the fit using leave-one-out (LOO) cross-validation (dashed lines).

**Table 2:** Intention-to-treat analyses

|  | CR for each variable independently |  | CR incl. all variables |  | CR after ADM reduction incl. all variables |  |
| --- | --- | --- | --- | --- | --- | --- |
|  | coefficients | p value | coefficients | p value | coefficients | p value |
| **Demographics** |  |  |  |  |  |  |
| Age | 0.19 | 0.28 | -0.22 | 0.61 | -0.12 | 0.77 |
| Male sex, No. (%) | 0.03 | 0.87 | 0.23 | 0.29 | 0.18 | 0.38 |
| Intelligence<sup>b</sup> | 0.29 | 0.13 | 0.28 | 0.34 | 0.18 | 0.51 |
| Site Berlin, No. (%) | -0.04 | 0.84 | -0.37 | 0.16 | -0.19 | 0.41 |
| **Clinical predictors** |  |  |  |  |  |  |
| *Current symptoms* |  |  |  |  |  |  |
| Residual depression<sup>b</sup> | 0.19 | 0.37 | 0.08 | 0.79 | -0.03 | 0.92 |
| Residual anxiety<sup>b</sup> | 0.21 | 0.24 | -0.27 | 0.39 | -0.20 | 0.51 |
| Somatic pain<sup>b</sup> | 0.31 | 0.08 | 0.44 | 0.08 | 0.22 | 0.32 |
| General impairment<sup>b</sup> | 0.32 | 0.04 | 0.33 | 0.36 | 0.30 | 0.38 |
| *Clinical history* |  |  |  |  |  |  |
| Age of onset | 0.05 | 0.78 | 0.06 | 0.89 | 0.01 | 0.98 |
| Chronicity<sup>b</sup> | -0.01 | 0.94 | -0.29 | 0.32 | -0.22 | 0.40 |
| Severity<sup>b</sup> | -0.04 | 0.83 | -0.07 | 0.77 | 0.14 | 0.54 |
| Number of prior episodes | 0.18 | 0.21 | 0.13 | 0.80 | -0.05 | 0.92 |
| Severity factor<sup>c</sup> | 0.25 | 0.13 | 0.10 | 0.89 | 0.21 | 0.76 |
| Comorbidities<sup>b</sup> | -0.03 | 0.88 | -0.08 | 0.76 | -0.08 | 0.73 |
| *Treatment* |  |  |  |  |  |  |
| Treated by GP, No. (%) | 0.36 | 0.03 | 0.66 | 0.005 | 0.41 | 0.043 |
| Length of ADM intake | -0.25 | 0.22 | -0.06 | 0.84 | -0.07 | 0.80 |
| Medication load<sup>c</sup> | -0.25 | 0.22 | -0.15 | 0.51 | -0.03 | 0.88 |
| Psychotherapy<sup>c</sup> | 0.01 | 0.96 | 0.34 | 0.15 | 0.06 | 0.77 |

a) Unless stated otherwise, mean (SD) are shown; b) Determined as follows: intelligence: Mehrfachwahl Wortschatz Test (Lehr, 2005); residual depression: Inventory of Depressive Symptomatology-Clinician Rated (Rush *et al.*, 1996); residual anxiety: screening generalised anxiety disorder (GAD-7; Spitzer *et al.*, 2006); somatic pain: somatisation subscale of symptom checklist 90 (SCL-90; Derogatis and Cleary, 1977) general impairment: global severity index of SCL-90; chronicity: numbers of months sick within the last 5 years; severity: symptoms during the last episode; comorbidities: number of past and present psychiatric diagnoses c) Computation of the variables is described in the section S1.4. GP = general practitioner; ADM = antidepressant medication; CR= cox regression; FU = Follow-up period (up to six months from end of discontinuation)

### 4.3 Prediction of relapse

To ascertain whether these findings could inform clinical practice, we next assessed how well clinical variables were able to predict relapses. Individual predictions could only meaningfully be assessed on the complete-case data. The multiple linear regression with all variables included achieved an area under the curve (AUC) of 0.76 with a sensitivity of 0.87 and specificity of 0.53 at the best cut-off (Fig. 2B). The model explained 21% of the variance. Such a performance is suggestive of clinical utility. However, with 18 predictor variables for 84 outcomes, this model may have overfitted the data and therefore may not generalize to new data.

We first examined overfitting through regularization via an elastic net, which pushes regression weights towards zero except for those predictor variables with most predictive power (Zou and Hastie, 2005). A standard approach with elastic nets, namely setting λto be one standard error larger than the value minimizing deviance, resulted in all regression weights being set to zero. A less stringent regularization using the value of λthat minimized deviance resulted in a model with non-zero weights for five variables only (intelligence, somatic pain, general impairment, severity factor and treatment by GP; Tab. 1) with an AUC of 0.74, a specificity of 0.66 and a sensitivity of 0.76 at the best cut-off value (Fig. 2B). Thus, five variables may suffice to predict relapse. However, since this is a within-sample analysis, it is still not clear whether and how well this result would generalise.

To determine how the models’ performances might generalise to new incoming patients, we approximated out-of-sample predictive accuracy using leave-one-out cross-validation (LOOCV). Here, all individuals are left out once, and the ability to predict whether they relapse using parameter estimates obtained from the other individuals’ data is assessed. Doing this without regularisation yielded a balanced accuracy of 0.49. With regularisation, the balanced accuracy was 0.46. Neither prediction exceeded chance.

### 4.4 Discontinuation Effect

The impact of antidepressant discontinuation on symptoms was examined by comparing changes in symptoms between the two main assessments in individuals randomized to groups 1W2 and 12W (Fig. 1). Discontinuation resulted in changes in residual symptoms in all four domains, including anxiety (F(1,89)=6.55, p=0.012), depression (F(1,89)=1.46, p=0.001) and general impairment (F(1,89)=9.99, p=0.002; Fig. 3A-D and Tab. S2). Post-hoc tests corrected for multiple comparisons indicated that no difference between groups exist at MA1, but did at MA2 and that the changes were due to an increase in symptoms in the group that discontinued their ADMs. For somatic pain, the interaction effect only showed a trend towards a significant difference (F(1,89)=3.31, p=0.072), and post-hoc tests of change only survived FDR correction in the discontinuation group.

**Figure 3:**
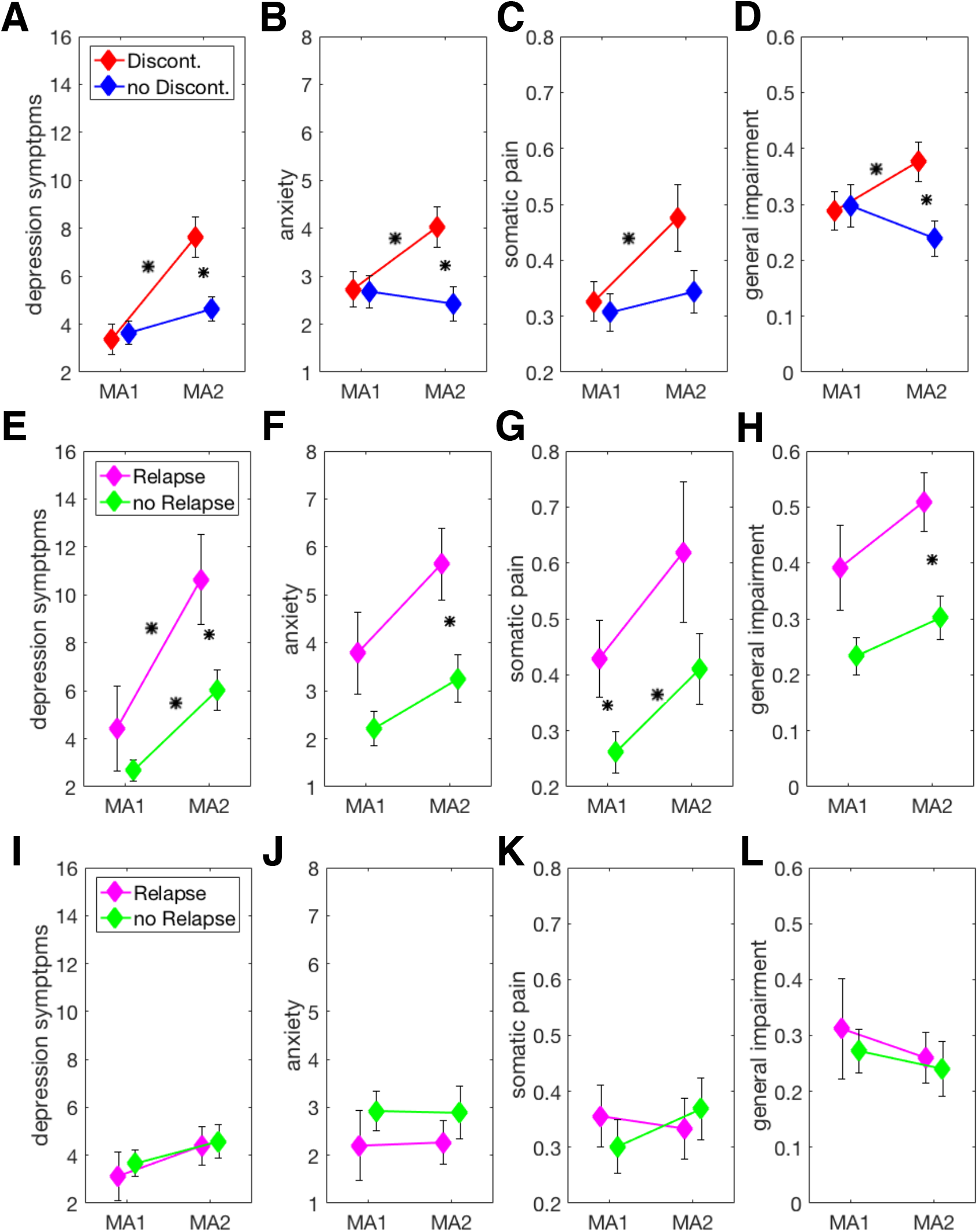
**(A-D) Discontinuation effects:** Changes in symptoms from main assessment one (MA1) to main assessment two (MA2) for depression (A), anxiety (B), somatic pain (C) and general impairment (D) in patients who discontinued between the two assessments and patients who did not discontinue. **(E-H) Discontinuation relapse interaction effects:** Changes in symptoms from MA1 to MA2 for depression (E), anxiety (F), somatic pain (G) and general impairment (H) in patients who discontinued and either relapsed or remained well during the follow-up period. **(I-L) Test-retest reliability for symptom measures:** Changes in symptoms from MA1 to MA2 for depression (I), anxiety (J), somatic pain (K) and general impairment (L) in patients who did not discontinue and either relapsed or remained well during the follow-up period. Asterisks indicate a significant difference at p < 0.05.

### 4.5 Association between discontinuation effect and relapse

We next asked whether the early effect of antidepressant discontinuation is associated with the ultimate risk of relapse. There was no interaction between the change in clinical measures before and after discontinuation (i.e. between the two main assessments in patients who discontinued before MA2, group 1W2; c.f. Fig. 1) and relapse (all p>0.05). Instead, the analysis revealed main effects of relapse across all domains (anxiety (F(1,40)=8.751, p = 0.005), general impairment (F(1,40)=11.001, p=0.003), depression (F(1,40)=5.615, p=0.023) and somatic pain (F(1,40)=4.709, p=0.036)). Relapsers in the group 1W2 had more symptoms before starting discontinuation and symptoms in both relapsers and non-relapsers increased after discontinuation to a similar extent (Fig. 3E-H), while there were no changes in the group that did not discontinue before MA2 (i.e. group 12W; Figure 3I-L, Table S2).

## 5 Discussion

Antidepressant medications are efficient in the prevention of relapses and relapse rates after discontinuation are high (Geddes *et al*., 2003). Relapse rates in our study were high, with one in three patients suffering a relapse within six months of discontinuation. This high relapse rate was observed even though the median duration of treatment was around two years, and hence at least as long as the duration of treatment recommended for recurrent illness (Bauer *et al*., 2013; NICE, 2019), and despite including only fully remitted patients with HAMD_17_ scores below 7. Relapses are not only important because they represent a period of renewed illness, but because any one episode has a 5-10% risk of becoming chronic (Hollon *et al*., 2006) and because early on in the disease additional episodes may mark the transition between those with a benign outcome and few lifetime episodes, and those with a malignant outcome and high risk of relapses (Keller *et al*., 1983, 1984; Monroe and Harkness, 2005, 2011). This situation makes it evident that there is a clinical need to establish biomarkers that can predict relapses specifically after antidepressant discontinuation, as such biomarkers could guide the discontinuation decision and in that way help reduce relapses and possibly even modify the long-term course of the illness.

A first pertinent step is the examination of the predictive power of clinical variables that are easily assessed in clinical practice. Our results suggest that such standard clinical variables carry at best weak predictive power. This conclusion relies on an examination of the likely generalisability of the associations. The approach is motivated by machine-learning approaches (Huys *et al*., 2016). Rather than asking how well a set of variables can predict a particular outcome *within* a given dataset, the prediction is assessed on out-of-sample data *not used* in ascertaining the prediction parameters. Such approaches are standard in the field of machine-learning, and are becoming more prominent in neuroscience and psychiatry (e.g.Chekroud *et al*., 2016; Wolfers *et al*., 2015; Dinga *et al*., 2018). We note that our cross-validation approach is not perfect as establishing a valid clinical predictor would ideally involve a fully independent dataset, but in our case this analysis indicates that the standard regression results do not carry predictive power.

Several aspects of the results from the standard approach are nevertheless noteworthy. First, in the full regression model and the intention-to-treat analyses including all predictors, only GP treatment emerged as significantly associated with relapse. This suggests that better treatment outcomes may be achieved when patients remain in specialist care, raising the question of what the active ingredient in this might be. Although in the current study psychotherapy did not appear to have an effect on relapse rates, our assessment of psychotherapeutic intervention strength was crude, and does leave room for the possibility that relapse risk could be mitigated by means of specific psychotherapeutic input. Indeed, psychotherapeutic techniques explicitly aimed at relapse have been developed (Hollon *et al*., 2005, 2014). Second, we did not replicate the effects of anxiety on relapse risk (Joliat *et al*., 2004), but the complete-case analyses replicated somatic pain as a risk factor (Fava *et al*., 2009). Third, the null findings do replicate null findings from RCTs (Berwian *et al*., 2017) in a naturalistic setting. Importantly, the two indicators which clinical guidelines emphasise, namely the number of prior episodes and the length of ADM treatment (Bauer *et al*., 2013; NICE, 2019), both failed to show an association with relapse risk in our naturalistic setting. This mirrors previous findings in RCTs (Berwian *et al*., 2017) and the consistent lack of coherent effects of these measures on relapse risk after ADM discontinuation suggests a revisiting of these recommendations. In a similar vein, we found no effect of residual symptoms, a decision criterion added in the newest version of the guidelines (NICE, 2019), on subsequent relapse risk. This is the case despite an influence of residual symptoms on overall relapse risk (Nierenberg *et al*., 2010) and symptom severity being the best predictor of disease course in studies using similar analyses approaches for patients in a depressive episode (Chekroud *et al*., 2016; Dinga *et al*., 2018). Finally, the lack of effects of any other clinical variable is still surprising given the relation to overall relapse risk of several of them as reviewed previously (Burcusa and Iacono, 2007; Hardeveld *et al*., 2010).

Next, discontinuation was associated with a robust increase in current symptoms across domains. Surprisingly, this increase in symptoms did not appear to be related to prospective relapses. The dissociation we observed raises the possibility that the mechanisms driving symptom increase after discontinuation differ from those driving subsequent relapse even though relapse trajectories on and off medication are similar (Gueorguieva *et al*., 2017). Clinically, the fact that transient symptomatic worsening does not relate to relapse may help clinicians and patients alike to hold their nerve in the face of early worsening of symptoms.

The study has strengths and limitations. Most prominently, the naturalistic setting of the study limits our ability to draw causal inferences: the pharmacological discontinuation effect is confounded with the potential psychological effect of knowing that the medication has been discontinued, and these cannot be disentangled. However, the naturalistic design increases the relevance for real-life outpatient care where these effects co-occur. A strength is the application of cross-validation to examine generalisability, but the small sample size is an important limitation. The small sample size also limits the identifiability of mechanistically heterogeneous subgroups. We also did not stratify for our randomisation variables when examining the effect of discontinuation. Finally, it might be that antidepressant discontinuation confers a vulnerability, but requires some additional stressor for a relapse to ensue. In that case, relapse after discontinuation would not be determined (and predictable) by characteristics before discontinuation, but by environmental influences thereafter.

### 5.1 Clinical implications

The results of the present study need to be replicated. Nevertheless, they are of potential clinical relevance and suggest several changes to the management of remitted depressive disorders. First, there may be a role for continued specialist care, in particular during and after the discontinuation phase. Second, prominent decision criteria currently used in clinical practice such as length of treatment, number of prior episodes and residual symptoms are poorly predictive of relapse, suggesting that guidelines for antidepressant discontinuation might have to be revisited. Third, both treatment providers and patients need to be informed that discontinuation may be accompanied by a transient re-emergence of depressive symptoms that do not necessarily indicate an imminent relapse.

### 5.2 Conclusion

Easily assessable demographic and clinical variables appear to be of limited use to guide antidepressant discontinuation decisions. Given the importance of the problem, more complex and costly measures should be evaluated.

## 5.3 Acknowledgements

This study was principally funded by a Swiss National Science Foundation grant (320030L_153449 / 1) to QJMH and by a Deutsche Forschungsgemeinschaft grant (WA 1539/5-1) to HW. Additional funds were provided by the Clinical Research Priority Program “Molecular Imaging” at the University of Zurich and by the René and Susanne Braginsky Foundation (to KES).

## 5.5 Contributions

QJMH and HW conceived and designed the study with critical input by KES and ES. IMB and JW collected the data under the supervision of QJMH and HW. KES was the study sponsor in Zurich. QJMH, ES, KES and HW acquired funding for the study. IMB planned and performed the analyses and wrote the manuscript under supervision by QJMH. All authors provided critical comments, read and approved the manuscript.

## 6 Supplementary Material

### S1 Supplementary Methods

#### S1.1 In- and Exclusion Criteria

Participants fulfilling the following inclusion criteria were eligible for participation in the study:

1. age 18-55 years
2. ability to consent and adhere to the study protocol
3. written informed consent
4. fluent in written and spoken German.

Patients had to additionally fulfil the following criteria:

1. currently under medical care with a psychiatrist or general practitioner for remitted Major Depressive Disorder and willing to remain in care for the duration of the study (approx. 9 months)
2. informed choice to discontinue medication (including willingness to taper the medication over at most 12 weeks) that was independent of study participation
3. clinical remission (HAMD_17_ of less than 7) had been achieved under therapy with Antidepressant Medication (ADM) without having undergone manualized psychotherapy; with no other concurrent psychotropic medication and had been maintained for a minimum of 30 days,
4. consent to information exchange between treating physician and study team members regarding inclusion/exclusion criteria and past medical history.

Any of the following exclusion criteria led to exclusion of an participant. This included the following general criteria

1. any disease of type and severity sufficient to influence the planned measurement or to interfere with the parameters of interest (This includes neurological, endocrinological, oncological comorbidities, a history of traumatic or other brain injury, neurosurgery or longer loss of consciousness.)
2. premenstrual syndrome (ICD-10 N94.3).

and MRI-related criteria

1. MRI-incompatible metal parts in the body,
2. inability to sit or lie still for a longer period,
3. possibility of presence of any metal fragments in the body,
4. pregnancy,
5. pacemaker, neurostimulator or any other head or heart implants,
6. claustrophobia and
7. dependence on hearing aid.

For patients the following additional criteria would led to exclusion:

1. current psychotropic medication other than antidepressants,
2. questionable history of major depressive episodes without complicating factors,
3. current acute suicidality,
4. lifetime or current axis II diagnosis of borderline or antisocial personality disorder,
5. lifetime or current psychotic disorder of any kind, bipolar disorder,
6. current posttraumatic stress disorder, obsessive compulsive disorder, or eating disorder
7. current drug use disorder (with the exception of nicotine) or within the past 5 years.

Healthy controls were excluded if there was a lifetime history of Diagnostic and Statistical Manual of Mental Disorders (4th ed., text rev.; DSM-IV-TR;American Psychiatric Association, 2000) axis I or axis II disorder with the exception of nicotine dependence.

#### S1.2 Procedures

Details for all interviews and questionnaires relevant for the present study are described in section S1.3.

##### S1.2.1 Recruitment

Participants were recruited using two types of strategies: 1) we informed outpatient clinics, practicing psychiatrist and general practitioners about the study with help of presentations and letters and distributed leaflets requesting them to inform potentially eligible patients about the option to participate in the study; 2) we recruited patients directly from the general population sending emails to staff and students of the universities, distributing leaflets to households, advertising in newspapers, streetcars and Google, and publicity about the study in newspaper articles. It might be of note for researcher planning future studies with a similar population, that we recruited a large majority of patients via the second strategy.

##### S1.2.2 Randomisation

After inclusion, participants were openly randomised into one of the two study arms, i.e. they either discontinued before the second main assessment (MA2; group 1W2) or did both MAs before discontinuation (group 12W). We assumed *a priori* that severity might affect relapse rates. Hence, to ensure that both groups would contain a sample with equal distributions of severity, we stratified patients into a severe and a non-severe group. If patients fulfilled either of the following criteria, they were assigned to the severe group: 1) more than three prior episodes, 2) more than seven depressive symptoms during the last episode, 3) severely impaired social functions, i.e. strongly isolated, disabled or aggressive, 4) engagement in activities almost nonexistent, 5) capacity to work almost nonexistent. We, additionally, stratified for site. Group membership was allocated by a randomisation algorithm. The first ten subjects at each site were not randomised, but all assigned to arm 1W2.

##### S1.2.3 Baseline Assessment

Participants who passed an initial telephone screening were invited for an on-site baseline assessment (BA).The first part of the BA was designed to assess if participants were eligible for the study. Inclusion criteria are listed in section S1.1. Participants underwent three clinical interviews to assess 1) current depressive symptoms, 2) axis-I disorders and 3) axis-2 disorders, i.e. personality disorders. Furthermore, we conducted a life-long medication screening on pharmaceuticals, a questionnaire to check for eligibility to enter a magnetic resonance imaging scanner, as well as individual questions relating to treatment. Participants had to sign that they discontinue their medication voluntarily, be in treatment with a physician who agreed to accompany the discontinuation process and wave the medical privilege of that physician with regard to the study team. If inclusion criteria were met, participants were randomised to either study arm, filled out a questionnaire batch assessing stable traits and underwent a short neuropsychological testing.

##### S1.2.4 Main Assessments

During both main assessments participants underwent four types of assessments: 1) an fMRI session including tasks to probe affective decision-making, automatic and voluntary emotion regulation, memory and a restingstate session, 2) a venipuncture to assess genetics (MA1 only), epigenetics, gene expression, cytokines, brain-derived neural growth factor, pharmacology level (before discontinuation only), C-reactive protein and a blood count, 3) behavioural testing including behavioural task to measure effort-related decision-making, impulsivity and emotion recognition and another questionnaire batch assessing state variables and 4) a questionnaire batch assessing state variables. In addition, current depressive symptoms were assessed in a clinical interview. In the two mornings before MA1, participants were asked to collect saliva tubes such that morning cortisol levels could be derived. Patients randomised to group 12W were asked to come for a third visit after discontinuation to undergo an additional venipuncture.

On top of these assessments, participants at the Zurich side were offered the option of additionally taking part in an electroenecelography (EEG) session including tasks measuring emotional reactivity, loudness dependent auditory potential, vigilance, resting-state and heartbeat evoked potentials.

##### S1.2.5 Follow-up Period and Final Assessment

During the follow-up period, patients were regularly contacted by phone and had to fill out short online questionnaire batches to assess symptom changes. Final assessment (FA) took either place if patients had a relapse or after six months. Current depressive symptoms were assessed and the state questionnaire batch was repeated.

#### S1.3 Questionnaires and Clinical Assessments

Clinical in- and exclusion criteria, as well as disease history and course, were assessed with the Structured Clinical Interview for DSM-IV (SCID) I and II to diagnose axis 1 disorders (major mental disorders) and axis II disorders (personality disorders), respectively (Wittchen and Fydrich, 1997). The Structured Interview Guide for Hamilton Depression Rating Scale (SIGH-D; Williams, 1988) consisting of 17 items was used to assess inclusion and the Inventory of Depressive Symptomatology Clinician Rated (IDS-C; Rush *et al*., 1996) with 30 items to quantify residual depression. Intelligence was assessed with the Mehrfachwahl Wortschatz Test (Lehr, 2005). The symptom checklist (SCL-90;Derogatis and Cleary, 1977) measures psychological impairment using nine subscales with 83 items in total and seven additional items. The global severity index (GSI) is the mean of all responses and is used to assess general psychological impairment. To assess somatic pain, we used the average score of somatisation subscale. This subscale contains twelve items and reflects simple somatic burden and functional disorder. We used a 7-item scale developed to screen for generalised anxiety disorder (GAD-7 Spitzer *et al*., 2006) to measure residual anxiety symptoms.

Additionally, we included questionnaires in the study but not in the reported analyses that tapped into the following domains: anhedonia/motivation, anxiety, mood, sexuality, discontinuation symptoms, personality, optimism/pessimism, emotion regulation, thinking style, resilience, self control, stressor load, quality of life, trauma, and demography. Questionnaires measuring stable traits and past experiences were assessed during BA. Measures assumed to differ within short time periods and be affected by discontinuation and disease state were assessed at MA1, MA2 and FA.

#### S1.4 Data Analysis

Disease severity corresponds to the first principal component of a principle component analysis including the variables number of past depressive episodes, age at illness onset, time in remission, time since depression onset, severity of last episode, time sick in total and time sick in the last five year as variables.

Medication load was based on the dose prior to discontinuation divided by the maximal allowed dose according to the Swiss compendium (www.compendium.ch) and by the weight of the participant.

Psychotherapy score was coded such that patients with no psychotherapy within the year before the study received a 0, patients reporting to have completed a psychotherapy within one year before the study a 0.5 and patients reporting to be in psychotherapy at the beginning of the study as 1. Significance was computed with a three-way chi-squared test.

For duration of antidepressant medication intake is the median and the interquartile range are reported. Significance was computed by means of the rank sum test.

### S2 Supplementary Results

Figure S1 shows reasons for dropouts in the patient sample.

Patients who did not complete the study had more residual symptoms (t(102) = −2.01, p = 0.047, CI = −3.74 - −0.02) and more symptoms during the last episode (t(102) = −2.09, p = 0.039, CI = −1.24 - −0.03) as shown in Table S1.

**Figure S1:**
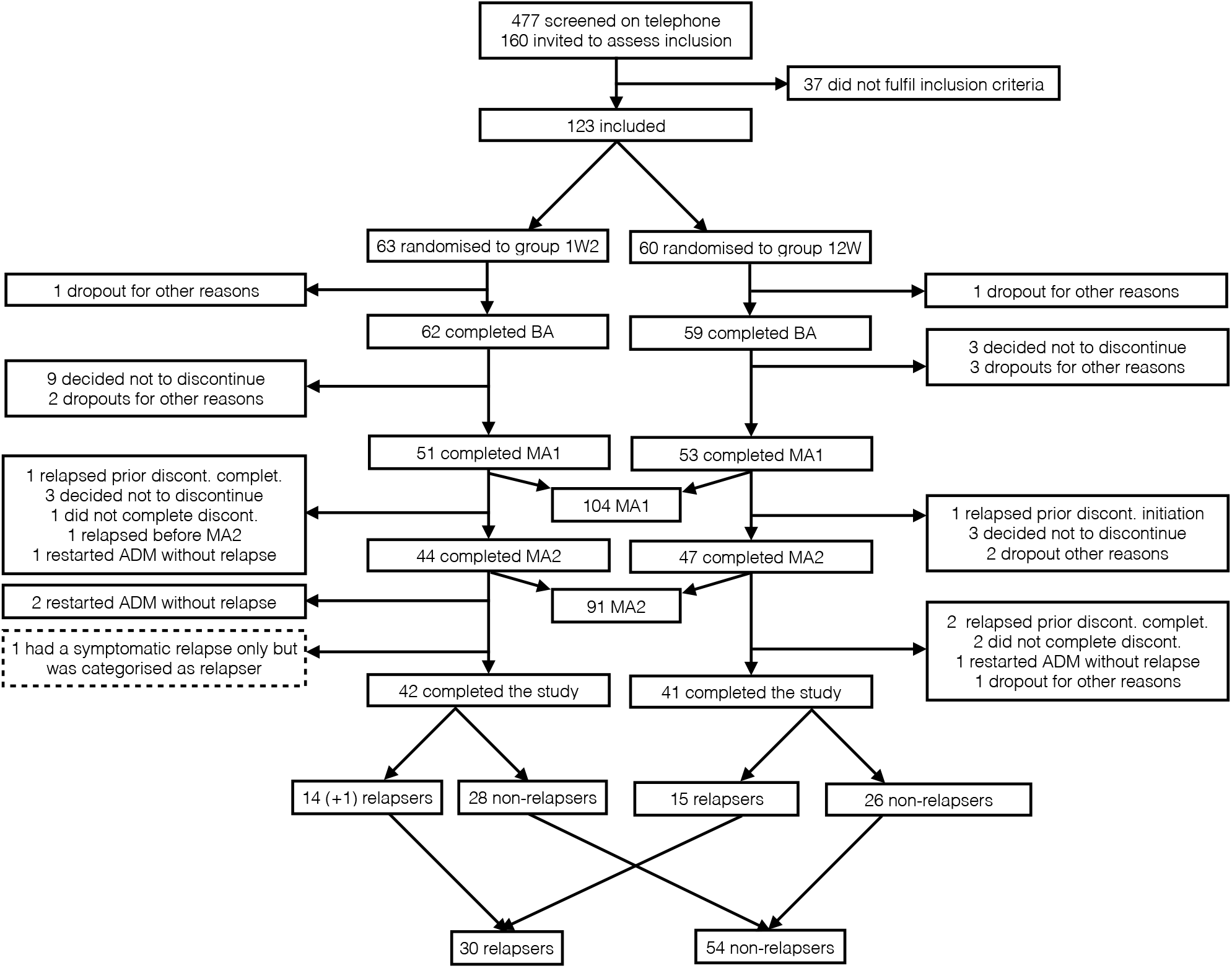
Consort Diagram: Depicted are reasons for dropouts and exclusion for patients throughout the study. (+ X) indicates the number of participants who relapsed after discontinuation but before MA2.

**Table S1:** Dropout Comparisons

|  | Study completed (n = 84) | Dropouts (n = 20) | p value |
| --- | --- | --- | --- |
| **Demographics** |  |  |  |
| Age | 34.79 (11.19) | 35.10 (11.02) | 0.91 |
| Male sex, No. (%) | 20 (24) | 4 (20) | 0.72 |
| Intelligence<sup>b</sup> | 28.40 (4.40) | 26.9 (4.49) | 0.17 |
| Site Berlin, No. (%) | 23 (27) | 5 (25) | 0.83 |
| **Clinical predictors** |  |  |  |
| *Current symptoms* |  |  |  |
| Residual depression<sup>b</sup> | 3.37 (3.73) | 5.25 (3.88) | 0.047 |
| Residual anxiety<sup>b</sup> | 2.73 (2.41) | 3.60 (1.90) | 0.13 |
| Somatic pain<sup>b</sup> | 0.32 (0.23) | 0.35 (0.24) | 0.63 |
| General impairment<sup>b</sup> | 0.29 (0.24) | 0.39 (0.25) | 0.09 |
| *Clinical history* |  |  |  |
| Age of onset | 24.23 (8.88) | 22.55 (9.60) | 0.46 |
| Chronicity<sup>b</sup> | 8.12 (9.73) | 10.00 (11.05) | 0.45 |
| Severity<sup>b</sup> | 7.01 (1.24) | 7.65 (1.18) | 0.039 |
| Number of prior episodes | 2.45 (1.61) | 2.50 (1.67) | 0.91 |
| Disease severity<sup>c</sup> | 0.02 (0.35) | 0.05 (0.48) | 0.74 |
| Comorbidities<sup>b</sup> | 0.76 (1.12) | 1.30 (1.26) | 0.06 |
| *Treatment* |  |  |  |
| Treated by GP, No. (%) | 18 (21) | 4 (20) | 0.89 |
| Length of ADM intake<sup>c</sup> | 23.5 (36) | 21.5 (22.5) | 0.16 |
| Medication load<sup>c</sup> | 0.01 (0.004) | 0.01 (0.004) | 0.32 |
| Psychotherapy<sup>c</sup> | 0.39 (0.39) | 0.38 (0.39) | 0.99 |

a) Unless stated otherwise, mean (SD) are shown; b) Determined as follows: intelligence: Mehrfachwahl Wortschatz Test (Lehr, 2005); residual depression: Inventory of Depressive Symptomatology-Clinician Rated (Rush *et al.*, 1996); residual anxiety: screening generalised anxiety disorder (GAD-7; Spitzer *et al.*, 2006); somatic pain: somatisation subscale of symptom checklist 90 (SCL-90; Derogatis and Cleary, 1977); general impairment: global severity index of SCL-90; chronicity: numbers of months sick within the last 5 years; severity: symptoms during the last episode; comorbidities: number of past and present psychiatric diagnoses c) Computation of the variables is described in the section S1.4. GP = general practitioner

Table S2 depicts details results of the discontinuation effect, the discontinuation relapse interaction effect and test-retest reliability for residual depression, residual anxiety, somatic pain and general impairment.

**Table S2:** Discontinuation effects

| Changes from MA1 to MA2 in all patients |  |  |  |  |  |  |  |  |
| --- | --- | --- | --- | --- | --- | --- | --- | --- |
|  | Discontinuation |  |  | No discontinuation |  |  | Discontinuation vs. No discont. |  |
|  | MA1 | MA2 | p value | MA1 | MA2 | p value | p value (MA1) | p value (MA2) |
| Residual depression | 3.39 (4.19) | 7.64 (5.69) | <0.001 | 3.64 (3.39) | 4.64 (3.40) | 0.11 | 0.75 | 0.006 |
| Residual anxiety | 2.73 (2.42) | 4.02 (2.81) | 0.009 | 2.68 (2.35) | 2.43 (2.40) | 0.73 | 0.93 | 0.009 |
| Somatic pain | 0.33 (0.23) | 0.48 (0.40) | 0.02 | 0.31 (0.22) | 0.34 (0.26) | 0.44 | 0.69 | 0.12 |
| General impairment | 0.29 (0.23) | 0.38 (0.23) | 0.03 | 0.30 (0.26) | 0.24 (0.22) | 0.10 | 0.85 | 0.02 |
| Changes from MA1 to MA2 in patients who discontinued |  |  |  |  |  |  |  |  |
|  | Relapsers |  |  | Non-relapsers |  |  | Relapsers vs. Non-relapsers |  |
|  | MA1 | MA2 | p value | MA1 | MA2 | p value | p value (MA1) | p value (MA2) |
| Residual depression | 4.43 (6.58) | 10.64 (7.07) | 0.014 | 2.68 (2.34) | 6.04 (4.44) | <0.001 | 0.21 | 0.02 |
| Residual anxiety | 3.79 (3.19) | 5.64 (2.79) | 0.09 | 2.21 (1.85) | 3.25 (2.61) | 0.07 | 0.07 | 0.04 |
| Somatic pain | 0.43 (0.26) | 0.62 (0.47) | 0.12 | 0.26 (0.20) | 0.41 (0.34) | 0.04 | 0.05 | 0.12 |
| General impairment | 0.39 (0.29) | 0.51 (0.20) | 0.14 | 0.23 (0.18) | 0.30 (0.20) | 0.11 | 0.07 | 0.01 |
| Changes from MA1 to MA2 in patients who did not discontinue |  |  |  |  |  |  |  |  |
|  | Relapsers |  |  | Non-relapsers |  |  | Relapsers vs. Non-relapsers |  |
|  | MA1 | MA2 | p value | MA1 | MA2 | p value | p value (MA1) | p value (MA2) |
| Residual depression | 3.13 (3.95) | 4.40 (3.11) | 0.60 | 3.65 (1.80) | 4.58 (3.6) | 0.60 | 0.83 | 0.88 |
| Residual anxiety | 2.20 (2.81) | 2.27 (1.75) | 0.95 | 2.92 (2.13) | 2.89 (2.81) | 0.95 | 0.89 | 0.89 |
| Somatic pain | 0.37 (0.21) | 0.33 (0.21) | 0.70 | 0.30 (0.25) | 0.37 (0.28) | 0.70 | 0.70 | 0.70 |
| General impairment | 0.31 (0.35) | 0.26 (0.17) | 0.79 | 0.27 (0.20) | 0.24 (0.25) | 0.79 | 0.79 | 0.79 |

Variables were measured with the following instruments: residual depression: Inventory of Depressive Symptomatology-Clinician Rated (Rush *et al.*, 1996); residual anxiety: screening generalised anxiety disorder (GAD-7; Spitzer *et al.*, 2006); somatic pain: somatisation subscale of symptom checklist 90 (SCL-90; Derogatis and Cleary, 1977); general impairment: global severity index of SCL-90. All p-values are corrected according to false-discovery-rate. MA1 = main assessment 1; MA2 = main assessment 2;

